# GLX10, a Novel Immunometabolic Modulator, Enhances Glycemic Control and Suppresses Inflammatory Signaling in a High-Fat Diet and Streptozotocin-Induced Rat Model of Type 2 Diabetes

**DOI:** 10.64898/2026.04.16.718956

**Authors:** Sherif Salah Abdul Aziz, Khalid F Kassem, Mohamed Sherif Salah

## Abstract

Type 2 diabetes mellitus (T2DM) is a progressive metabolic disorder characterized by persistent hyperglycemia, insulin resistance, and chronic low-grade inflammation. Despite the widespread use of established therapies such as metformin, long-term glycemic control remains suboptimal, and disease progression is often not adequately prevented. This highlights the need for novel therapeutic strategies that address both metabolic dysfunction and the underlying immunometabolic components of the disease. In this study, GLX10 (GLXM100) was evaluated as a novel immune modulator in a high-fat diet (HFD) and low-dose streptozotocin (STZ)-induced rat model of T2DM over a 91-day period. Glycemic outcomes were assessed using terminal random blood glucose and oral glucose tolerance testing (OGTT), with glucose exposure quantified by area under the curve (AUC 0-120). Complementary in vitro investigations were performed in hepatic and macrophage cell models to assess cytocompatibility, nitric oxide production, and modulation of pro-inflammatory cytokines, including IL-6 and TNF-α. GLX10 treatment resulted in a significant reduction in random blood glucose levels and a marked improvement in glucose tolerance compared to diabetic control animals. Importantly, GLX10 demonstrated greater improvement in OGTT AUC compared to metformin under the same experimental conditions, indicating enhanced dynamic glucose regulation. In vitro, GLX10 maintained viability in normal hepatic cells while significantly suppressing nitric oxide production and inflammatory cytokine outputs in macrophages, supporting a favorable safety and immune profile. Collectively, these findings demonstrate that GLX10 exerts robust antidiabetic activity through a dual mechanism involving metabolic regulation and suppression of inflammatory signaling. The integration of in vivo efficacy with supportive in vitro safety and mechanistic data provides a strong preclinical foundation and supports the further development of GLX10 as a promising therapeutic candidate for T2DM.

## INTRODUCTION

Type 2 diabetes mellitus is a chronic metabolic disorder characterized by persistent hyperglycemia resulting from a combination of insulin resistance and progressive pancreatic beta-cell dysfunction [1,2,24–27]. The global prevalence of this condition has increased markedly in recent decades, driven by changes in lifestyle, dietary patterns, and rising rates of obesity, making it a major public health concern worldwide [2]. Long-term hyperglycemia is associated with both microvascular and macrovascular complications, including nephropathy, neuropathy, retinopathy, and cardiovascular disease, which collectively contribute to increased morbidity and mortality [3].

Current pharmacological strategies are primarily directed toward reducing blood glucose levels and improving insulin sensitivity. First-line therapies such as metformin act mainly by suppressing hepatic glucose production and enhancing peripheral glucose uptake [4]. Although effective during early stages of the disease, many patients fail to maintain sustained metabolic control and often require treatment intensification as the condition progresses. In addition, several therapeutic options are limited by adverse effects, reduced long-term efficacy, or diminished responsiveness in advanced insulin-resistant states. These limitations highlight the need for approaches that address broader aspects of disease biology rather than focusing solely on glycemic control [5].

Experimental models that closely replicate human disease are essential for the evaluation of candidate therapies. The combination of high-fat diet feeding followed by low-dose streptozotocin administration represents a well-established model that reproduces key features of type 2 diabetes mellitus, including insulin resistance, partial pancreatic beta-cell impairment, and sustained hyperglycemia [6,7,31–33]. This model allows assessment of both baseline glycemic status and dynamic glucose handling through oral glucose tolerance testing, providing a reliable framework for evaluating therapeutic responses.

Beyond altered glucose metabolism, type 2 diabetes mellitus is also associated with chronic low-grade inflammation. Elevated levels of pro-inflammatory mediators, including tumor necrosis factor alpha and interleukin-6, interfere with insulin signaling pathways and contribute to metabolic dysfunction [8–10,28–30]. Activation of immune cells, particularly macrophages, leads to increased production of nitric oxide and other inflammatory mediators, linking immune responses with metabolic regulation. These observations support the development of therapeutic strategies that incorporate modulation of inflammation-related pathways.

Cell-based experimental systems provide complementary insight into safety and mechanistic direction. Hepatic cell models are widely used to evaluate cytotoxicity and metabolic responses, reflecting the central role of the liver in glucose regulation and xenobiotic metabolism. Macrophage-derived cells are commonly employed to study inflammatory signaling, nitric oxide production, and cytokine responses. In addition, cellular processes such as apoptosis, cell cycle progression, and autophagy serve as indicators of how candidate compounds influence cellular stress responses and metabolic adaptation.

The investigational compound evaluated in this study has been proposed to influence both metabolic and immune-related processes relevant to type 2 diabetes mellitus. Rigorous evaluation using validated in vivo disease models and standardized cellular assays is required to establish its therapeutic potential and safety profile. In the present work, the compound was assessed in a high-fat diet and streptozotocin-induced rat model, with emphasis on glycemic control and glucose tolerance, alongside complementary analyses of cytotoxicity, inflammatory mediator modulation, nitric oxide production, and signaling pathway activity in hepatic and immune cell systems.

The proposed mechanism of action is based on a coordinated regulatory framework that targets cellular adaptation rather than direct glucose lowering. This approach involves controlled activation of innate immune pathways together with downstream mitochondrial and transcriptional responses. The complementary peptide component, administered subcutaneously, is designed to induce a controlled intracellular stress signal through interaction with pattern recognition receptors, leading to activation of adaptive cellular programs such as the mitochondrial unfolded protein response and mitophagy, which are essential for maintaining mitochondrial integrity.

In parallel, the intramuscular component functions as an immunomodulatory signal that enhances adaptive immune responsiveness while contributing to reduction of chronic inflammation. This includes modulation of macrophage activity and improvement of the cellular environment associated with metabolic stress. Together, these processes may influence transcriptional regulation and contribute to sustained changes in gene expression patterns linked to metabolic homeostasis.

At the cellular level, these coordinated responses converge on improved mitochondrial function, including enhanced turnover of damaged organelles, stabilization of mitochondrial membrane potential, reduction of reactive oxygen species, and improved energy production. These effects may alleviate metabolic stress and contribute indirectly to improved insulin sensitivity and systemic glucose regulation.

From a pharmacodynamic perspective, the compound represents a coordinated system in which immune activation and immunomodulation act together to engage endogenous cellular repair mechanisms. Rather than targeting a single molecular pathway, this framework suggests a system-level mode of action involving interaction between immune signaling, mitochondrial dynamics, and regulatory processes controlling cellular adaptation under chronic metabolic stress.

While this mechanistic framework is supported by preclinical observations, it should be considered a working hypothesis that requires further experimental validation, particularly with respect to causal relationships between immune activation, mitochondrial function, and long-term metabolic outcomes.

### AIM AND OBJECTIVES

Aim: The primary aim of this study was to evaluate the antidiabetic efficacy and preclinical safety profile of GLX10 using an integrated in vivo high-fat diet and low-dose streptozotocin rat model of type 2 diabetes mellitus, supported by in vitro assays assessing cytocompatibility and immunometabolic responses in relevant hepatic and macrophage cell lines.

Objectives: The specific objectives of this study were:

In-vivo objectives:

1. To assess the effect of GLX10 on weekly random blood glucose (RBS) and body weight trends in a high-fat diet + STZ rat model.
2. To evaluate glucose handling at the end of the study using oral glucose tolerance test (OGTT) and AUC□–□□□ analysis.
3. To compare the glycemic effects of GLX10-only treatment with metformin-only treatment, while also evaluating the sequential metformin→GLX10 regimen as an exploratory treatment approach.
4. To evaluate systemic safety and organ function using terminal biochemical markers including ALT, AST, creatinine, and BUN.
5. To evaluate hematological safety using CBC and differential leukocyte count.
6. To assess organ-level effects through organ weights (liver, pancreas, kidney) and histological assessments, including liver and kidney histopathology and pancreatic insulin staining.

In-vitro objectives:

7 To determine the cytotoxicity profile of GLX10 in hepatic cell lines (BNL-1S and HepG2) using SRB viability assays and IC□□ estimation.
8 To investigate the effect of GLX10 on inflammatory mediator output in macrophages (RAW 264.7), including nitric oxide inhibition and modulation of key pro-inflammatory cytokines (IL-6 and TNF-α).
9 To evaluate the impact of GLX10 on cellular fate and stress-response pathways including apoptosis, autophagy, and cell cycle distribution in relevant in-vitro models.
10 To assess the influence of GLX10 on macrophage signaling pathways relevant to inflammation and metabolism (including NF-κB/mTOR axis and SAMHD1 phosphorylation) as supportive mechanistic evidence.

## Materials and Methods

This study consisted of two integrated components designed to provide both efficacy and mechanistic insight into the activity of GLX10. The first component was an in vivo rat study aimed at evaluating the metabolic and organ-level effects of GLX10 in a high-fat diet and streptozotocin (HFD–STZ) model of diabetes as shown in **Supplementary Figure S1**. The second component comprised an in vitro package performed by NAWAH Scientific (Cairo, Egypt), which assessed cytotoxicity, inflammatory responses, cell death modalities, and pathway signaling relevant to the interpretation of the in vivo findings. Although the overall program initially began as a broader dose-ranging exploration involving 120 male Wistar rats distributed across 20 experimental groups, the present report was intentionally restricted to the six most informative groups, representing the key experimental comparisons and the GLX10 regimen(s) that produced biologically interpretable responses as showed in supplementary S1.

Accordingly, the final reported dataset included 36 rats in total, with 6 rats per group. Male Wistar rats were housed under controlled laboratory conditions at 22 ± 2°C, 40 ± 5% humidity, and a 12 h light/12 h dark cycle, with lights on at 07:00. Animals were group-housed at three rats per cage in polycarbonate cages containing Aspen wood-chip bedding and had ad libitum access to food and water. All animals underwent a 7-day acclimatization period before study initiation.

To induce insulin resistance prior to chemical induction of diabetes, animals received a high-fat diet formulated from standard rodent chow supplemented with vegetable margarine. The final dietary composition provided approximately 45% of energy from fat, 35% from carbohydrates, and 20% from protein. This diet was administered ad libitum for four weeks before streptozotocin exposure and was continued throughout the study period. Diabetes was induced using freshly prepared streptozotocin dissolved in 0.1 M citrate buffer at pH 4.5. Because streptozotocin is light sensitive and unstable in solution, it was prepared under reduced light, kept chilled at approximately 4°C, and administered within 10 minutes of preparation.

Animals were fasted for 6 h before induction, then streptozotocin was administered intraperitoneally at 25 mg/kg. To reduce the risk of acute hypoglycemia following injection, animals received 10% glucose solution for 24 h after streptozotocin administration. Diabetes induction was confirmed after 72 h using a fasting blood glucose threshold of greater than 250 mg/dL. The present report includes six predefined in vivo groups: therapeutic GLX10, sequential metformin followed by GLX10, metformin control, normal control, diabetic control, and prophylactic GLX10. GLX10 was administered subcutaneously at a dose of 5 units per rat twice daily, whereas metformin was administered orally at 250 mg/kg.

The study timeline included acclimatization, a 4-week HFD induction phase, streptozotocin injection on Day 28, diabetes confirmation during Days 29–35, treatment phase 1 from Days 36–57, treatment phase 2 from Days 70–90 where applicable, weekly monitoring of body weight and random blood glucose, and oral glucose tolerance testing on Day 91, followed by terminal blood collection and organ harvest.

Body weight was recorded weekly using a calibrated digital balance, and random blood glucose was measured weekly using tail-vein sampling with sterile single-use lancets and a validated glucometer. Sampling was performed as consistently as possible within the same weekly time window to minimize diurnal variability.

On Day 91, animals underwent an oral glucose tolerance test after a 6 h fast. Glucose was administered orally at 2 g/kg, and blood glucose measurements were obtained at 0, 15, 30, 60, and 120 min. OGTT results were analyzed both as individual time-point curves and as the total area under the curve from 0 to 120 min. At study termination, animals were euthanized using carbon dioxide inhalation with gradual fill, followed by a secondary physical confirmation method, in accordance with facility-approved humane euthanasia procedures.

Terminal blood was collected immediately after confirmation of death and divided into serum tubes for clinical chemistry analyses, including ALT, AST, creatinine, and BUN, and EDTA tubes for complete blood count and differential leukocyte analysis. Serum samples were allowed to clot, centrifuged, and stored under cold-chain conditions until analysis. Following blood collection, the liver, pancreas, and kidneys were excised, weighed, grossly inspected, and processed for histological evaluation. Liver and kidney tissues were prepared for histopathology, while pancreatic tissue was prepared for insulin immunostaining.

Histological samples were fixed in 10% neutral buffered formalin, processed through routine dehydration and paraffin embedding, sectioned at 3–5 µm, and stained with hematoxylin and eosin. Pancreatic insulin staining was performed using a validated anti-insulin primary antibody with standard HRP/DAB detection and hematoxylin counterstaining.

The in vitro component was performed by NAWAH Scientific and was restricted in this project to the comparison between control and GLX10 at the GLX M100 concentration. Additional compound codes or groups reported by the external laboratory were not considered in the present analysis. For cell-based assays involving BNL cells and flow cytometry packages, cells were maintained in DMEM supplemented with 10% heat-inactivated fetal bovine serum, penicillin (100 U/mL), and streptomycin (100 µg/mL), and incubated at 37°C in a humidified 5% CO□ atmosphere.

Cytotoxicity in BNL and HepG2 cells was evaluated using the sulforhodamine B assay, a colorimetric method that estimates cell mass based on total cellular protein content [19,20]. Cells were seeded in 96-well plates, allowed to attach, and then exposed to a concentration series of GLX10. After the designated exposure period, cells were fixed with trichloroacetic acid, stained with SRB dye, washed to remove excess stain, and the bound dye was solubilized for absorbance measurement using a plate reader. Cell viability was calculated relative to untreated controls, and IC□□ values were derived using nonlinear dose–response curve fitting. Nitric oxide production in RAW 264.7 macrophages was quantified indirectly by measuring nitrite accumulation in culture supernatants using the Griess reaction [23,43–45].

Following treatment under inflammatory stimulation conditions, supernatants were mixed with Griess reagent, incubated in the dark at room temperature, and absorbance was measured at 540 nm. Nitrite concentrations were determined using a standard calibration curve, and relative changes versus control were calculated accordingly.

Inflammatory cytokine outputs were assessed using sandwich ELISA kits for IL-6 and TNF-α. The assays were performed according to standard ELISA methodology using pre-coated plates, sample and standard incubation, biotinylated detection antibodies, avidin-HRP conjugate, substrate development, and absorbance reading at 450 nm. IL-6 and TNF-α kits from Elabscience were used, and plate reading was performed using a BMG Labtech FLUO star Omega instrument. Flow cytometry studies in BNL cells were conducted to evaluate apoptosis, autophagy, and cell cycle distribution.

Apoptosis was analyzed by Annexin V-FITC/propidium iodide staining, allowing discrimination of live, early apoptotic, late apoptotic, and necrotic cells [22]. Autophagy-related readouts were assessed using acridine orange staining, which accumulates in acidic vesicular organelles [21]. Cell cycle analysis was performed after ethanol fixation followed by RNase A and propidium iodide staining to determine DNA-content distribution across G0/G1, S, and G2/M phases.

Flow acquisition was carried out on an ACEA NovoCyte flow cytometer and analysis was performed using NoVo Express software, with approximately 12,000 events recorded per sample.

To explore pathway-level effects, RAW 264.7 macrophages were cultured in DMEM supplemented with 10% heat-inactivated fetal bovine serum and antibiotics, then seeded in 6-well plates and stimulated with lipopolysaccharide at 1 μg/mL to induce inflammation. GLX10 was applied at non-cytotoxic concentrations established from SRB assays, with untreated and LPS-only groups serving as controls.

Western blot analysis was then performed to evaluate total SAMHD1, phospho-SAMHD1 (Thr592), mTOR pathway markers including phospho-mTOR, phospho-S6, and phospho-4EBP1, and NF-κB pathway markers including phospho-p65, total p65, and IκBα. Protein samples were extracted using standard lysis procedures, quantified, separated by SDS-PAGE, transferred to membranes, probed with primary and HRP-conjugated secondary antibodies, and detected by chemiluminescence. Densitometric values were normalized to housekeeping controls and reported as fold change relative to control

### Statistical Analysis

All quantitative data generated from both in vivo and in vitro experiments were analyzed using appropriate statistical methods to ensure robustness and reproducibility of the findings. Data are presented as mean ± standard error of the mean (SEM), unless otherwise specified. For comparisons involving multiple experimental groups, statistical significance was evaluated using analysis of variance (ANOVA), followed by post hoc multiple comparison testing where applicable. In cases where group variances were unequal or sample sizes were limited, Welch’s correction was applied to improve statistical reliability [34–37].

To control for type I error arising from multiple comparisons, adjusted p-values were calculated using Holm’s correction method. Statistical significance was defined at a threshold of p < 0.05 after adjustment. For OGTT analysis, glucose excursion was quantified using the area under the curve (AUC□–□□□), calculated by the trapezoidal rule to provide an integrated measure of glucose exposure over time. In vitro dose–response relationships, including IC□□ estimation for cytotoxicity and nitric oxide inhibition assays, were determined using nonlinear regression models based on a four-parameter logistic (4PL) equation. All statistical analyses were performed using validated statistical software packages, and calculations were cross-checked to ensure consistency with experimental datasets. Detailed supporting analyses are provided in the Supplementary Materials.

## Results

In Vivo Glycemic Control and Glucose Tolerance, to evaluate the metabolic efficacy of GLX10, both longitudinal glycemic monitoring and dynamic glucose challenge testing were performed in the high-fat diet/streptozotocin (HFD/STZ) model of type 2 diabetes.

Longitudinal random blood glucose (RBG) measurements demonstrated a clear divergence between experimental groups following diabetes induction. Diabetic Control animals exhibited sustained hyperglycemia throughout the study period, whereas the GLX10 therapeutic group (D36–57) showed a progressive and sustained reduction in glucose levels, approaching near-physiological values at late time points. This sustained effect is illustrated in the longitudinal glucose profile (**Figure 2**), confirming a durable improvement in chronic glycemic burden. To further assess glucose handling capacity, an oral glucose tolerance test (OGTT) was conducted at Day 91. The Normal Control group demonstrated efficient glucose clearance, with a low peak and rapid return toward baseline levels. In contrast, Diabetic Control animals showed markedly impaired glucose tolerance, with sustained hyperglycemia throughout the 120-minute observation period.

Notably, the GLX10-treated group exhibited a substantially improved glucose excursion profile, characterized by a lower peak and faster decline toward baseline compared to Diabetic Control (**Figure 1A**). This improvement was quantitatively supported by a marked reduction in total glycemic exposure, with OGTT AUC□–□□□ reduced by approximately 42% relative to Diabetic Control (32,972.50 vs 57,021.25 mg·min/dL) (**Figure 1B**). In addition, the 120-minute glucose level in the GLX10 group (198.17 mg/dL) was significantly lower than that of Diabetic Control (346.17 mg/dL), indicating improved late-phase glucose clearance. In contrast, other intervention arms, including Metformin-treated and prophylactic GLX10 groups, exhibited severe post-load hyperglycemia, with values reaching the assay ceiling (600 mg/dL at 60 min), reflecting persistent impairment in glucose tolerance [14,38,39].

**Figure 1A.**
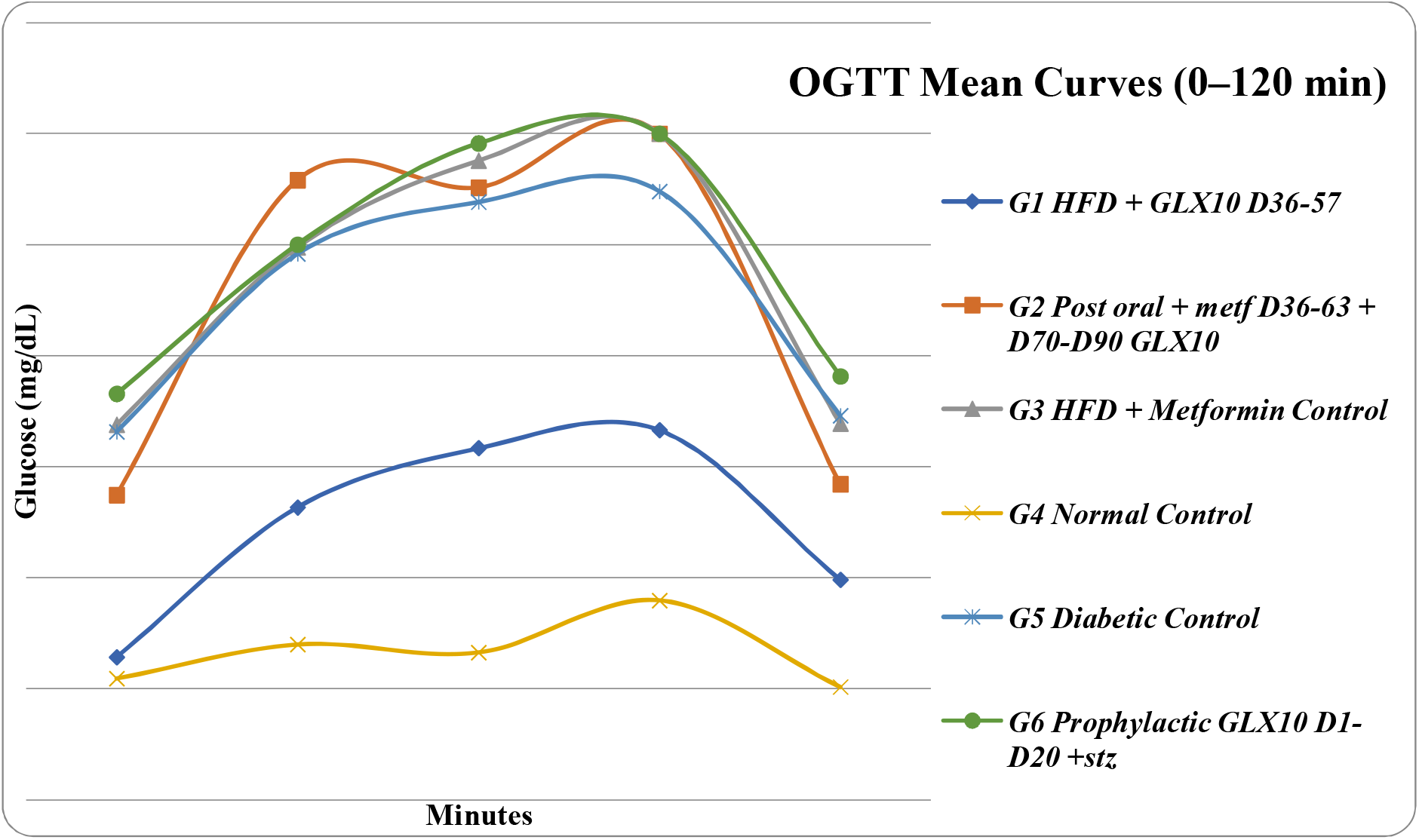
OGTT baseline and post-load glycemic response at D91.

**Figure 1A:**
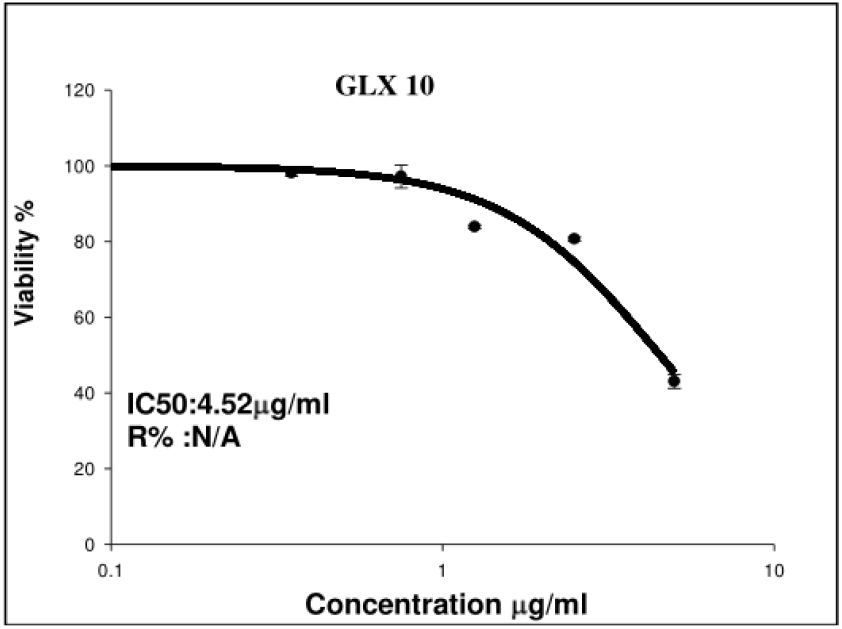
SRB dose-response viability in BNL cells (72 h). Dose-dependent reduction in BNL viability following 72 h exposure to GLX10 using SRB protein-staining readout.

**Figure 1B:**
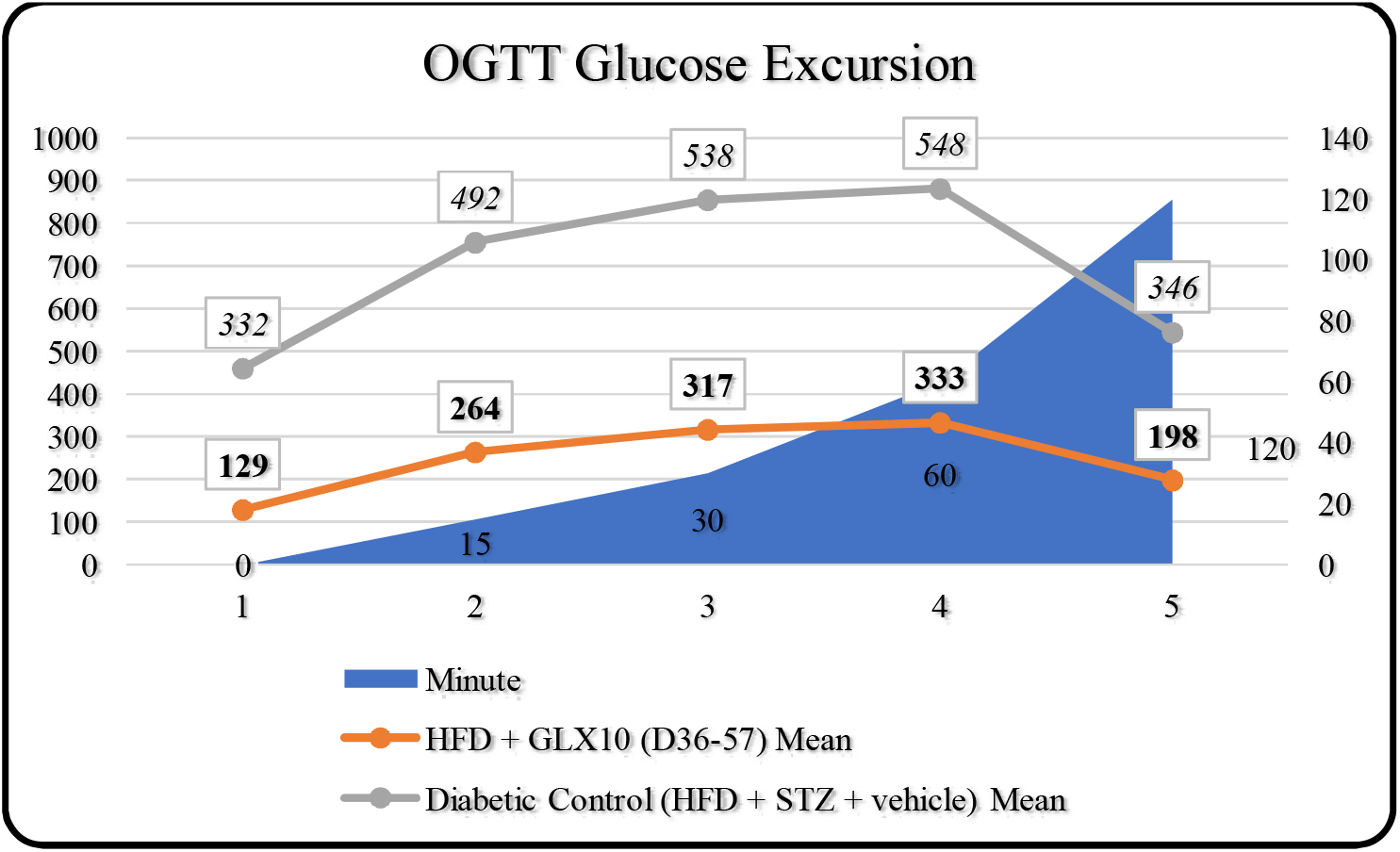
OGTT glucose excursion curve diabetic vs normal rat AUC comparison.

**Figure 2.**
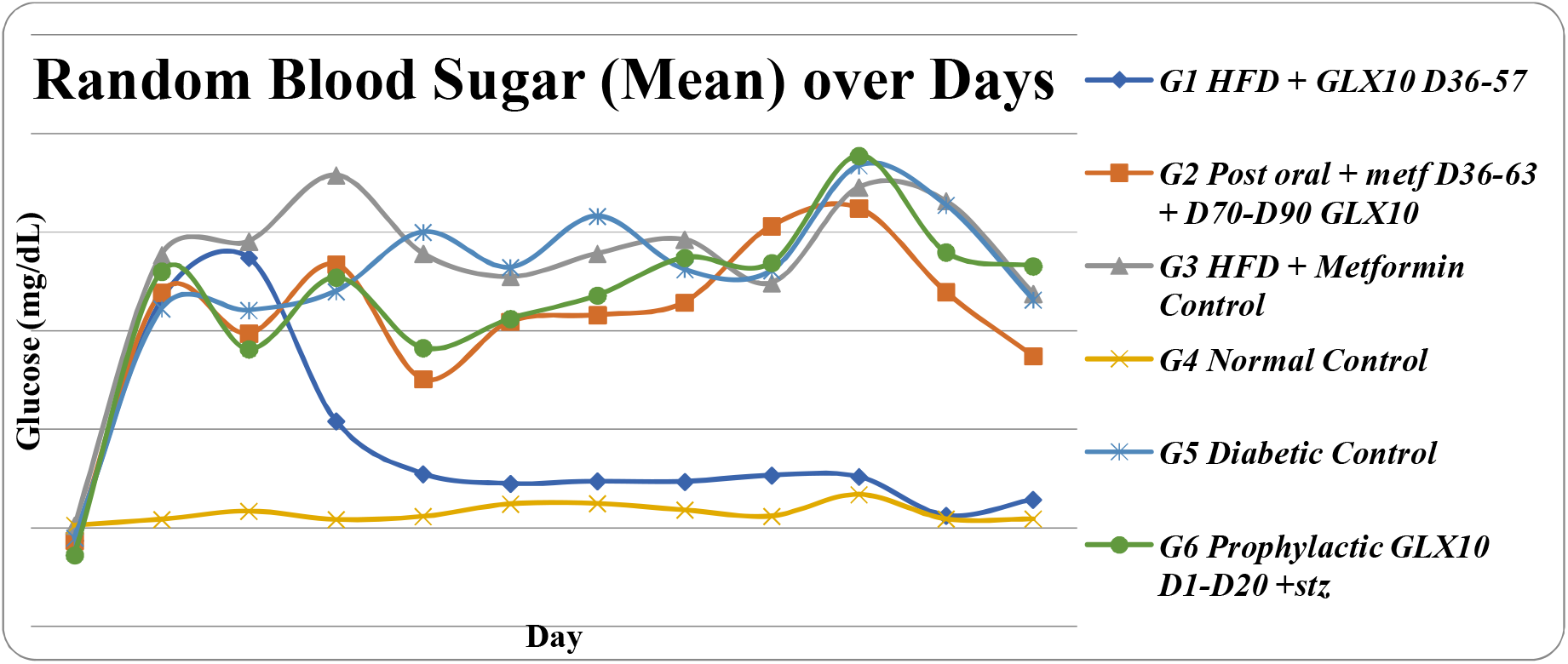
Random blood glucose (non-fasting) trends across the study timeline.

**Figure 2.**
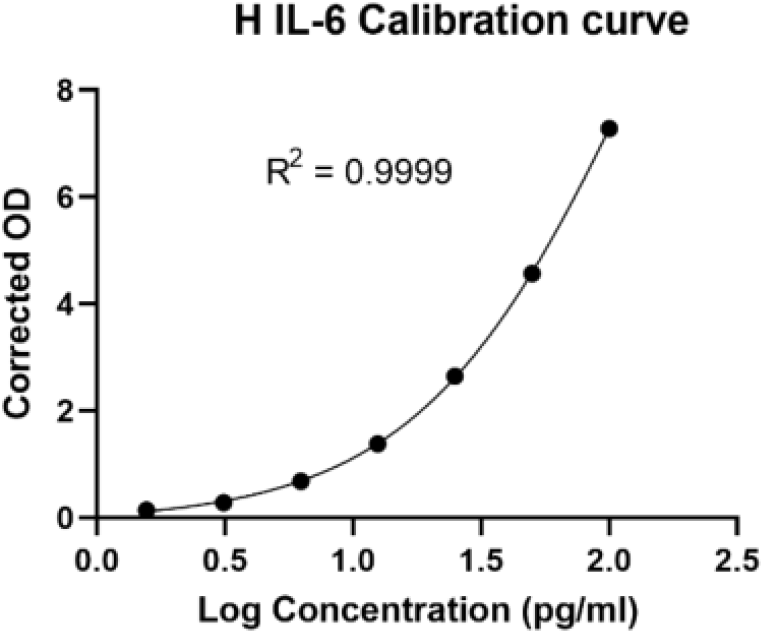
Human IL-6 ELISA readout (Control vs GLX10 conditions; reported in pg/mL).

Statistical analysis using Welch’s t-test with Holm correction confirmed that GLX10 (D36–57) was the only intervention achieving significant reductions in both RBG at Day 91 and OGTT AUC□–□□□ compared to Diabetic Control. These effects were associated with very large effect sizes (RBG: d = −3.752; OGTT AUC: d = −3.358), supporting a robust therapeutic impact.

**In Vitro Cytotoxicity and Selectivity Profile**, the cytotoxic and cytocompatibility profile of GLX10 was assessed using the sulforhodamine B (SRB) assay across three biologically distinct cell lines: BNL (normal hepatocytes), HepG2 (hepatocellular carcinoma), and RAW 264.7 (macrophages). GLX10 demonstrated differential sensitivity between normal and malignant cells. In HepG2 cells, viability decreased more sharply with increasing concentrations compared to BNL cells, indicating enhanced susceptibility of the carcinoma line (**Figure 3**). This observation was supported by IC□□ values derived from dose–response analysis, showing a lower IC□□ in HepG2 (≈ 2.68 µg/mL) relative to BNL cells (≈ 4.52 µg/mL), consistent with selective cytotoxicity toward malignant cells.

**Figure 3B:**
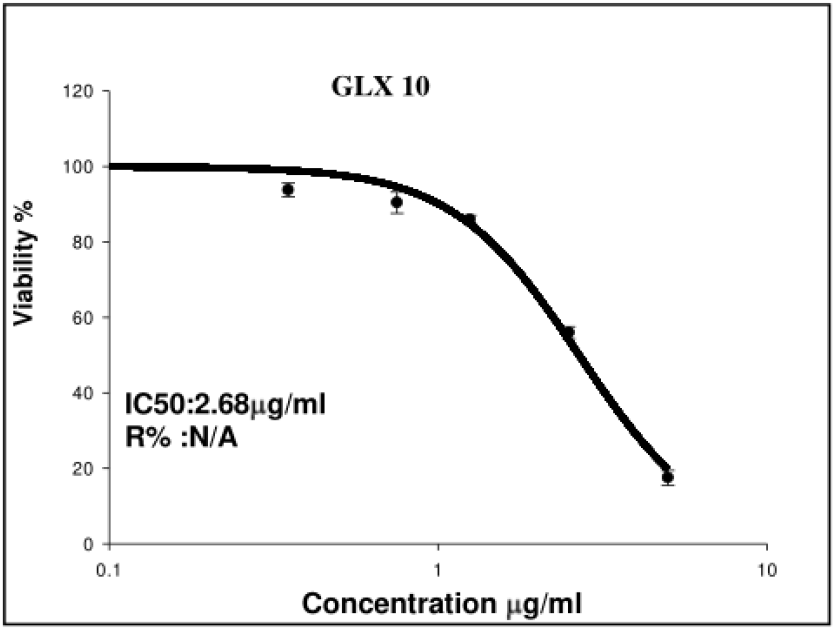
SRB dose-response viability in HepG2 cells (72 h). GLX10 induces stronger viability suppression in HepG2 carcinoma than in BNL normal liver across matched concentrations, consistent with higher sensitivity of the carcinoma model in SRB analysis.

In contrast, RAW 264.7 macrophages maintained relatively high viability across the tested concentration range, with no SRB IC□□ reached within the experimental window. This indicates that macrophage viability is largely preserved under conditions where GLX10 exerts functional effects, supporting a non-cytotoxic mechanism in immune cells.

Anti-inflammatory Activity in Macrophage Model, the anti-inflammatory activity of GLX10 was evaluated in RAW 264.7 macrophages through measurement of nitric oxide (NO) production using the Griess reaction [8,9]. GLX10 produced a clear concentration-dependent inhibition of NO production [10,15], increasing from approximately 18.46% at the lowest concentration (0.19 µg/mL) to 73.17% at the highest tested concentration (2.50 µg/mL), indicating strong suppression of inflammatory activation (**Figure 4 A**). Importantly, this effect occurred in the absence of significant macrophage cytotoxicity, supporting a true immunomodulatory mechanism rather than a non-specific toxic effect.

**Figure 4A:**
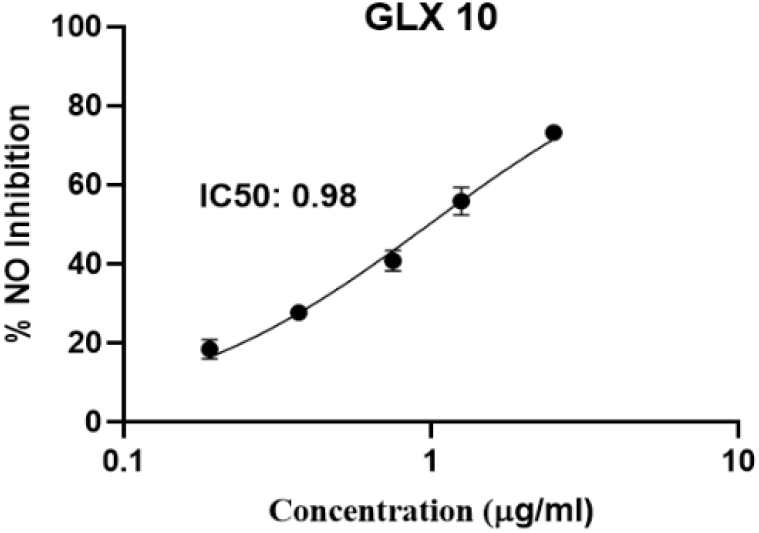
Dose-response inhibition of nitric oxide production in RAW 264.7 cells by GLX10 at 48 h. Dose-dependent inhibition of nitric oxide production (nitrite readout quantified via the Griess reaction) by GLX10 in RAW 264.7 macrophages at 48 h, reaching approximately 73% inhibition at 2.5 µg/mL.

**Figure 4C:**
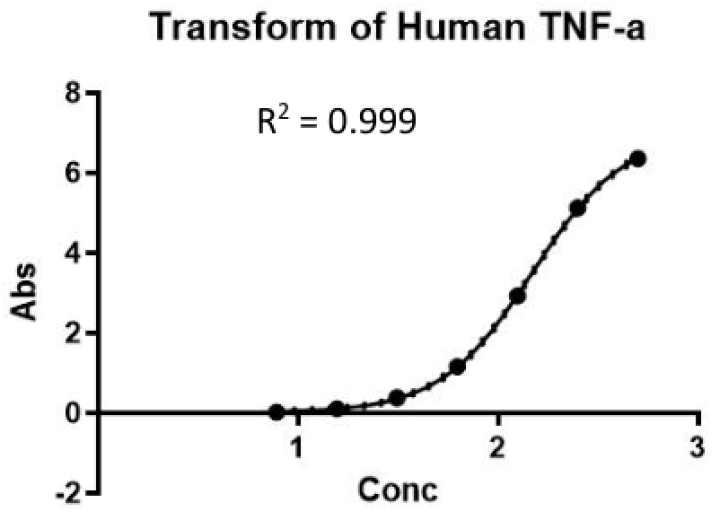
Human TNF-α ELISA readout (Control vs GLX10 conditions; reported in pg/mL).

This anti-inflammatory effect was further supported by cytokine analysis. GLX10 treatment resulted in reduced levels of key pro-inflammatory mediators, including IL-6 and TNF-α, compared to control conditions, indicating coordinated suppression of inflammatory signaling pathways (**Figure 4 B, C**).

**Cellular Mechanisms: Apoptosis, Autophagy**, and Cell Cycle Flow cytometry analysis in BNL cells revealed significant alterations in cellular state following GLX10 exposure. Apoptosis profiling demonstrated a marked redistribution of cell populations [13], with a reduction in the dominant viable quadrant and a corresponding increase in apoptotic fractions (**Supplementary Figure S 4**). Total cell death increased substantially from approximately 0.98% in control conditions to 20.86% following GLX10 treatment indicating activation of cell death pathways. Autophagy-related analysis using acridine orange staining showed increased fluorescence intensity metrics (Mean X and Median X), suggesting enhanced autophagic activity (**Supplementary Figure S 5**), although the gated population percentage (P1%) was reduced under GLX10 conditions Cell cycle analysis further demonstrated a shift in cell distribution, characterized by a decrease in G1 phase and a marked increase in Sub-G1 fraction (3.38% → 26.76%), indicative of DNA fragmentation and apoptotic progression (**Supplementary Figure S 6**).

**Signaling Pathway Modulation, western** blot analysis in RAW 264.7 cells revealed modulation of key signaling pathways involved in inflammation and cellular metabolism. GLX10 treatment resulted in reduced phosphorylation of mTOR pathway components (phospho-mTOR, phospho-S6, phospho-4EBP1) and decreased phospho-NF-κB p65 levels, while total NF-κB p65 remained relatively unchanged and IκBα expression was increased (**Supplementary Figure S 7**). In parallel, GLX10 altered SAMHD1 signaling, with increased total SAMHD1 expression and reduced phosphorylation at Thr592 suggesting modulation of immune-regulatory pathways.

Overall, the complete dataset demonstrates that GLX10 produces a consistent and robust improvement in glycemic control in vivo, while simultaneously exerting anti-inflammatory, cytomodulatory, and signaling effects in vitro. The convergence of these findings across all figures supports a coherent immune mechanism underlying the observed therapeutic activity. Comprehensive supporting data, including full tabulated datasets, extended statistical analyses, and additional experimental results, are provided in the Supplementary Materials (Figures S1– S10) to maintain clarity and conciseness of the main manuscript.

## Discussion

Type 2 diabetes mellitus is increasingly recognized as a complex disorder characterized not only by persistent hyperglycemia and insulin resistance, but also by chronic low-grade inflammation that contributes to disease progression. Pro-inflammatory mediators such as tumor necrosis factor alpha and interleukin-6 impair insulin signaling pathways, while activation of transcriptional regulators including nuclear factor kappa B further sustains metabolic dysfunction. Although metformin remains a central therapeutic option, its primary effects are largely limited to metabolic regulation and do not fully address the inflammatory processes that contribute to insulin resistance in advanced disease states [8,9].

In the present study, the investigational compound evaluated in this work was associated with a marked improvement in systemic glycemic control in a validated high-fat diet and streptozotocin-induced rat model [14]. Therapeutic administration resulted in a clear reduction in terminal random blood glucose levels at Day 91, indicating a decrease in chronic glycemic burden under non-fasting conditions. This effect remained statistically significant following multiple comparison correction, supporting the reliability of the observed findings. Improvement in glucose tolerance was further supported by oral glucose tolerance testing, where treatment reduced the total glucose exposure over a one-hundred-and-twenty-minute period compared with diabetic control animals. Under the same experimental conditions, the observed effect was greater than that achieved with metformin, although this comparison should be interpreted within the limitations of the experimental model.

This observation is relevant because the area under the oral glucose tolerance curve reflects dynamic glucose handling and provides a broader assessment of metabolic regulation than single time-point measurements. However, interpretation of these findings should consider that several experimental groups reached assay ceiling values, which may have reduced the apparent magnitude of intergroup differences and potentially masked partial improvements in more severe phenotypes.

Findings from cellular experiments provided additional insight into the biological effects observed in vivo. The compound inhibited nitric oxide production in macrophage-derived cells, indicating suppression of immune cell activation [10,15]. In addition, treatment reduced the production of pro-inflammatory mediators, including tumor necrosis factor alpha and interleukin 6, both of which are associated with impaired insulin signaling and metabolic dysfunction. These observations support the presence of an anti-inflammatory effect at the cellular level.

At the signaling level, treatment was associated with reduced activation of nuclear factor kappa B and the mechanistic target of rapamycin pathway, which are central regulators of inflammatory signaling and cellular metabolic responses [16,17,40–42]. Nuclear factor kappa B regulates transcription of inflammatory genes, whereas the mechanistic target of rapamycin pathway integrates nutrient sensing with cellular growth and energy balance. The observed modulation of these pathways suggests that the compound influences interconnected regulatory networks linking immune signaling and metabolic control. In addition, changes in SAMHD1 expression and phosphorylation status indicate a possible role in immune regulation, although the functional implications remain to be clarified [18].

The combined in vivo and cellular findings support a coherent biological interpretation. The reduction in glycemic burden and improvement in glucose tolerance are consistent with attenuation of inflammation-associated insulin resistance. This pattern supports the interpretation that the compound may act through coordinated regulation of immune and metabolic processes rather than functioning solely as a direct glucose-lowering intervention [46–51]. Interestingly, the therapeutic regimen produced the most pronounced effects, whereas the prophylactic regimen did not result in comparable improvements. This observation suggests a stage-dependent activity, where the compound may exert greater effects in established disease states characterized by active inflammatory signaling. Such a pattern is consistent with current understanding that interventions targeting inflammation tend to be more effective when pathological pathways are already active.

Despite these promising findings, several limitations should be considered. The study is preclinical in nature, and extrapolation to human physiology requires further validation. In addition, although modulation of key signaling pathways was observed, the causal relationships and upstream regulatory mechanisms remain to be established through targeted experimental studies.

Overall, the findings indicate that the investigational compound improves systemic glucose regulation while influencing inflammatory signaling pathways. This integrated effect distinguishes it from conventional therapeutic approaches and supports further investigation into its potential for translational development in type 2 diabetes mellitus.

## Conclusion

In this study, therapeutic administration of the investigational compound evaluated in this work was associated with a marked improvement in glycemic control within a high-fat diet and streptozotocin-induced rat model of type 2 diabetes mellitus. Treatment resulted in a substantial reduction in total glucose exposure during oral glucose tolerance testing over a one-hundred-and-twenty-minute period when compared with diabetic control animals, indicating improved regulation of glucose handling under the conditions of this model.

Complementary findings from cellular experiments demonstrated a consistent biological profile characterized by reduced production of nitric oxide and pro-inflammatory mediators, including tumor necrosis factor alpha and interleukin 6, together with modulation of key regulatory pathways involved in inflammation and cellular metabolism. These observations support an integrated effect on both metabolic regulation and inflammation-associated processes.

Taken together, the findings suggest that the investigational compound improves systemic glucose regulation while influencing pathways linked to immune and metabolic function. Although these results are consistent with a coordinated biological mechanism, further studies are required to establish causal relationships and to determine the relevance of these effects in clinical settings.

## Supporting information

https://doi.org/10.5281/zenodo.19610307

## Future Research Directions

Future studies should focus on expanding cohort size and refining dose–response evaluation to strengthen statistical power and confirmatory interpretation. Direct assessment of insulin sensitivity and secretion (e.g., insulin tolerance testing or clamp-based approaches) will be essential to clarify the underlying mechanism of glycemic improvement. In addition, tissue-level validation of signaling pathways in metabolically relevant organs (liver, skeletal muscle, and adipose tissue) is required to confirm the proposed NF-κB and mTOR modulation in vivo.

Long-term studies are also necessary to evaluate durability of response and extended safety, given the chronic nature of T2DM therapy. Finally, pharmacokinetic and pharmacodynamic (PK/PD) characterization will be critical to support dose optimization and translational progression toward clinical development.

### Credit Author Statement

Sherif Salah and Mohamed Sherif contributed equally to the conceptualization of the study, development of the scientific framework, experimental design, execution of the experimental work, data acquisition, and preparation of the original manuscript draft. Khalid Kassem contributed to critical review of the manuscript, provided medical and scientific guidance, and supported the interpretation of the findings.

## Ethical Considerations

All in vivo experimental procedures were conducted in accordance with internationally recognized guidelines for the care and use of laboratory animals. The study design, animal handling, and experimental procedures were aligned with institutional animal care and use committee (IACUC) principles and complied with the recommendations outlined in the Guide for the Care and Use of Laboratory Animals and ARRIVE reporting guidelines. Animals were housed under controlled environmental conditions and monitored regularly to ensure welfare throughout the study period. All procedures involving induction of diabetes, blood sampling, and treatment administration were performed using standardized protocols designed to minimize stress and variability. At study termination, euthanasia was carried out using carbon dioxide inhalation with gradual fill, followed by a secondary physical confirmation method, in accordance with the American Veterinary Medical Association (AVMA) guidelines for humane euthanasia. All efforts were made to reduce animal suffering and to use the minimum number of animals necessary to achieve scientifically valid results.

### Confidentiality Statement

This manuscript contains data related to the investigational candidate GLX10. Certain methodological details, including specific formulation and dosing parameters, are described at a general level due to their proprietary nature. Full technical information may be made available upon reasonable request and subject to appropriate agreements.

## Supplementary Materials

The Supplementary Materials include additional figures (Figures S1–S9) providing extended experimental data, detailed analyses, and supporting results complementary to the main findings.

The datasets and supplementary materials supporting this study are available at: https://doi.org/10.5281/zenodo.19610307

## REFERENCES

[1] DeFronzo RA, Ferrannini E, Zimmet P, Alberti KGMM. International Textbook of Diabetes Mellitus. 4th ed. Wiley-Blackwell; 2015.

[2] International Diabetes Federation. IDF Diabetes Atlas. 10th ed. Brussels; 2021.

[3] Forbes JM, Cooper ME. Mechanisms of diabetic complications. Physiol Rev. 2013;93(1):137–188.

[4] Rena G, Hardie DG, Pearson ER. The mechanisms of action of metformin. Diabetologia. 2017;60(9):1577–1585.

[5] Taylor R. Type 2 diabetes: etiology and reversibility. Diabetes Care. 2013;36(4):1047–1055.

[6] Srinivasan K, Viswanad B, Asrat L, Kaul CL, Ramarao P. Combination of high-fat diet and low-dose streptozotocin-treated rat as a model for type 2 diabetes. Pharmacol Res. 2005;52(4):313–320.

[7] Skovsø S. Modeling type 2 diabetes in rats. J Diabetes Investig. 2014;5(4):349–358.

[8] Hotamisligil GS. Inflammation and metabolic disorders. Nature. 2006;444:860–867.

[9] Donath MY, Shoelson SE. Type 2 diabetes as an inflammatory disease. Nat Rev Immunol. 2011;11:98–107.

[10] Nathan C, Ding A. Nonresolving inflammation. Cell. 2010;140(6):871–882.

[11] Donato MT, Tolosa L, Gómez-Lechón MJ. Human hepatocyte models. Methods Mol Biol. 2015;1250:77–93.

[12] Park J, et al. Macrophage activation pathways. Nat Commun. 2014;5:3264.

[13] Mizushima N, Levine B. Autophagy mechanisms. Nat Cell Biol. 2010;12:823–830.

[14] Ayala JE, et al. Metabolic testing standards. Dis Model Mech. 2010;3(9–10):525–534.

[15] Dandona P, Aljada A, Bandyopadhyay A. Inflammation, and insulin resistance. Trends Immunol. 2004;25(1):4–7.

[16] Liu T, Zhang L, Joo D, Sun SC. NF-κB signaling. Signal Transduct Target Ther. 2017;2:17023.

[17] Saxton RA, Sabatini DM. mTOR signaling. Cell. 2017;168(6):960–976.

[18] Cheung PHH, et al. SAMHD1 in immune regulation. Trends Immunol. 2025.

[19] Skehan P, et al. SRB assay. J Natl Cancer Inst. 1990;82(13):1107–1112.

[20] Vichai V, Kirtikara K. SRB method. Nat Protoc. 2006;1(3):1112–1116.

[21] Thomé MP, et al. Acridine orange staining. Biotechniques. 2016.

[22] Bio-Techne. Annexin V Apoptosis Detection Protocol.

[23] Sigma-Aldrich. Griess Reagent Protocol.

[24] American Diabetes Association. Standards of Medical Care in Diabetes—2024. Diabetes Care. 2024;47(Suppl 1):S1–S350.

[25] World Health Organization. Global report on diabetes. Geneva: WHO; 2024.

[26] GBD 2021 Diabetes Collaborators. Global, regional, and national burden of diabetes, 1990–2021. Lancet. 2023.

[27] International Diabetes Federation. IDF Diabetes Atlas update. Brussels; 2024.

[28] Weisberg SP, McCann D, Desai M, Rosenbaum M, Leibel RL, Ferrante AW. Obesity is associated with macrophage accumulation in adipose tissue. J Clin Invest. 2003;112(12):1796–1808.

[29] Xu H, Barnes GT, Yang Q, Tan G, Yang D, Chou CJ, et al. Chronic inflammation in fat plays a crucial role in the development of obesity-related insulin resistance. J Clin Invest. 2003;112(12):1821–1830.

[30] Shoelson SE, Lee J, Goldfine AB. Inflammation and insulin resistance. J Clin Invest. 2006;116(7):1793–1801.

[31] Lenzen S. The mechanisms of alloxan- and streptozotocin-induced diabetes. Diabetologia. 2008;51(2):216–226.

[32] Reed MJ, Meszaros K, Entes LJ, Claypool MD, Pinkett JG, Gadbois TM, et al. A new rat model of type 2 diabetes: the fat-fed, streptozotocin-treated rat. Metabolism. 2000;49(11):1390–1394.

[33] Skovsø S, et al. Modeling type 2 diabetes in rodents using high-fat diet and STZ combinations. Diabetology. 2020.

[34] Welch BL. The generalization of Student’s problem when several different population variances are involved. Biometrika. 1947;34(1–2):28–35.

[35] Holm S. A simple sequentially rejective multiple test procedure. Scand J Stat. 1979;6:65–70.

[36] Benjamini Y, Hochberg Y. Controlling the false discovery rate: a practical approach. J R Stat Soc B. 1995;57(1):289–300.

[37] Shapiro SS, Wilk MB. An analysis of variance test for normality. Biometrika. 1965;52(3–4):591–611.

[38] Andrikopoulos S, Blair AR, Deluca N, Fam BC, Proietto J. Evaluating the glucose tolerance test in mice. Am J Physiol Endocrinol Metab. 2008;295:E1323–E1332.

[39] Pruessner JC, Kirschbaum C, Meinlschmid G, Hellhammer DH. Two formulas for AUC calculation. Psychoneuroendocrinology. 2003;28(7):916–931.

[40] Hayden MS, Ghosh S. Shared principles in NF-κB signaling. Cell. 2008;132(3):344–362.

[41] Kim J, Kundu M, Viollet B, Guan KL. AMPK and mTOR signaling in autophagy regulation. Nat Cell Biol. 2011;13(2):132–141.

[42] Li X, et al. Emerging roles of mTOR and NF-κB signaling in metabolic diseases. Cell Metab. 2023.

[43] Green LC, Wagner DA, Glogowski J, Skipper PL, Wishnok JS, Tannenbaum SR. Analysis of nitrate and nitrite. Anal Biochem. 1982;126(1):131–138.

[44] Carvalho-Filho MA, Ueno M, Hirabara SM, Seabra AB, Carvalheira JB, de Oliveira MG, et al. S-nitrosation of insulin receptor signaling proteins. Diabetes. 2005;54(4):959–967.

[45] Pilon G, Penicaud L, Poirier P, et al. Role of nitric oxide in metabolic regulation. J Physiol. 2010.

[46] Kowluru RA, et al. Metabolic memory and epigenetics in diabetes. Diabetes. 2018.

[47] Pirola L, Balcerczyk A, Okabe J, El-Osta A. Epigenetic phenomena in diabetes complications. Nat Rev Endocrinol. 2015;11(4):220–228.

[48] Sun G, et al. Epigenetic regulation of metabolic genes. J Biol Chem. 2007.

[49] Yoshizaki T, et al. Epigenetic control in metabolic disease. Endocrinology. 2009.

[50] Yuan T, et al. Mitochondrial epigenetics in metabolic disorders. Trends Endocrinol Metab. 2022.

[51] Feng X, et al. Emerging role of mitochondrial epigenetics in diabetes. Cell Metab. 2025.

