## Supplementary material for "GLX10, a Novel Immunometabolic Modulator, Enhances Glycemic Control and Suppresses Inflammatory Signaling in a High-Fat Diet and Streptozotocin-Induced Rat Model of Type 2 Diabetes": https://doi.org/10.5281/zenodo.19610307

GLX10 Figures and Supplementary Dataset

Figure 1 A.

OGTT Mean Curves (0–120 min)

|  | G1 HFD + GLX10 D36-57 | G2 Post oral + metf D36-63 + D70-D90 GLX10 | G3 HFD + Metformin Control | G4 Normal Control | G5 Diabetic Control | G6 Prophylactic GLX10 D1-D20 +stz |
| --- | --- | --- | --- | --- | --- | --- |
| 0 | 128.667 | 274.5 | 337.833 | 109.167 | 331.5 | 365.667 |
| 15 | 263.5 | 558.167 | 497.667 | 139.833 | 491.833 | 500.167 |
| 30 | 316.667 | 551.5 | 575.667 | 132.667 | 538.333 | 591.333 |
| 60 | 333 | 600 | 600 | 179.5 | 548 | 600 |
| 120 | 198.167 | 284.333 | 339.167 | 101.667 | 346.167 | 381.5 |

Figure 1 A: OGTT baseline and post-load glycemic response at D91.

Figure 1 B.

OGTT Glucose Excursion

|  | Minute | HFD + GLX10 (D36-57) Mean | Diabetic Control (HFD + STZ + vehicle) Mean |
| --- | --- | --- | --- |
| 1 | 0 | 128.667 | 331.5 |
| 2 | 15 | 263.5 | 491.833 |
| 3 | 30 | 316.667 | 538.333 |
| 4 | 60 | 333 | 548 |
| 5 | 120 | 198.167 | 346.167 |

Figure 1B: OGTT glucose excursion curve diabetic vs normal rat AUC comparison.

Figure 2.

Random Blood Sugar (Mean) over Days

|  | G1 HFD + GLX10 D36-57 | G2 Post oral + metf D36-63 + D70-D90 GLX10 | G3 HFD + Metformin Control | G4 Normal Control | G5 Diabetic Control | G6 Prophylactic GLX10 D1-D20 +stz |
| --- | --- | --- | --- | --- | --- | --- |
| 21 | 87.5 | 87.1667 | 102.5 | 102.833 | 89.8333 | 72.5 |
| 24 | 331.5 | 339.167 | 377.333 | 108.833 | 322.333 | 360.333 |
| 28 | 374 | 297.333 | 390.667 | 117.167 | 320.833 | 281.5 |
| 35 | 208.667 | 367.5 | 458.167 | 108.5 | 340 | 354.333 |
| 42 | 155 | 251.167 | 378.5 | 111.667 | 400.333 | 282.667 |
| 49 | 145 | 309.5 | 355 | 124.5 | 364.333 | 312.333 |
| 56 | 147.333 | 316.167 | 378.667 | 124.833 | 416.333 | 336 |
| 63 | 147 | 329 | 392.667 | 118.333 | 362.333 | 374.167 |
| 70 | 153.333 | 406.333 | 348.667 | 112.167 | 361.333 | 369 |
| 77 | 151.833 | 424.5 | 445.833 | 134.167 | 467.833 | 477.667 |
| 84 | 112.667 | 339.5 | 431.833 | 109.167 | 427.667 | 379.833 |
| 91 | 128.667 | 274.5 | 337.833 | 109.167 | 331.5 | 365.667 |

Figure 2: Random blood glucose (non-fasting) trends across the study timeline.

Figure 3.

| 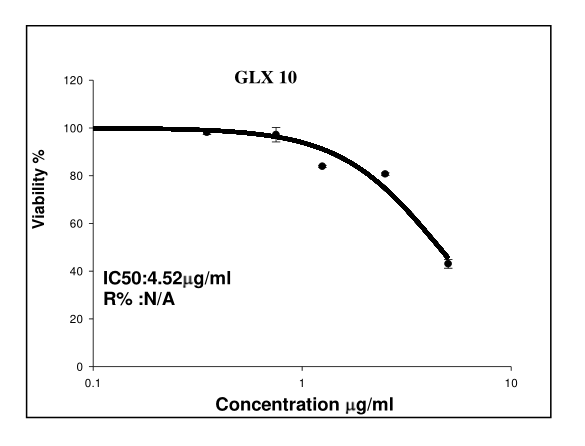 | 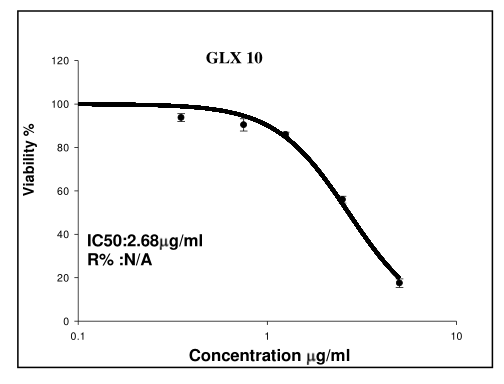 |
| --- | --- |
| Figure 3 A: SRB dose-response viability in BNL cells (72 h). Dose-dependent reduction in BNL viability following 72 h exposure to GLX10 using SRB protein-staining readout. | Figure 3 B: SRB dose-response viability in HepG2 cells (72 h). GLX10 induces stronger viability suppression in HepG2 carcinoma than in BNL normal liver across matched concentrations, consistent with higher sensitivity of the carcinoma model in SRB analysis. |

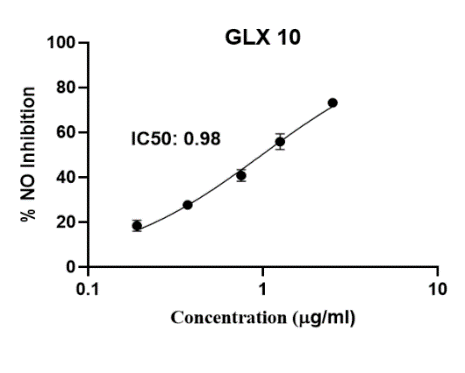

Figure *4 A*: Dose-response inhibition of nitric oxide production in RAW 264.7 cells by GLX10 at 48 h. Dose-dependent inhibition of nitric oxide production (nitrite readout quantified via the Griess reaction) by GLX10 in RAW 264.7 macrophages at 48 h, reaching approximately 73% inhibition at 2.5 µg/mL.

Figure 4

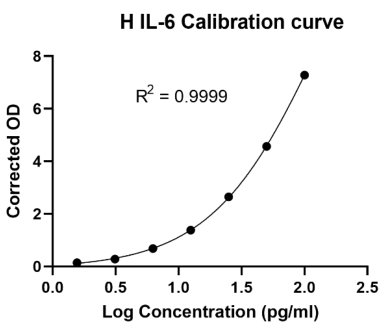

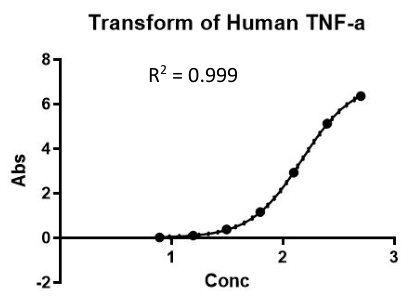

Figure 4 C: Human TNF-α ELISA readout

(Control vs GLX10 conditions; reported in pg/mL).

Figure 4 B: Human IL-6 ELISA readout

(Control vs GLX10 conditions; reported in pg/mL).

Supplementary FigureS1 Experimental Design.png

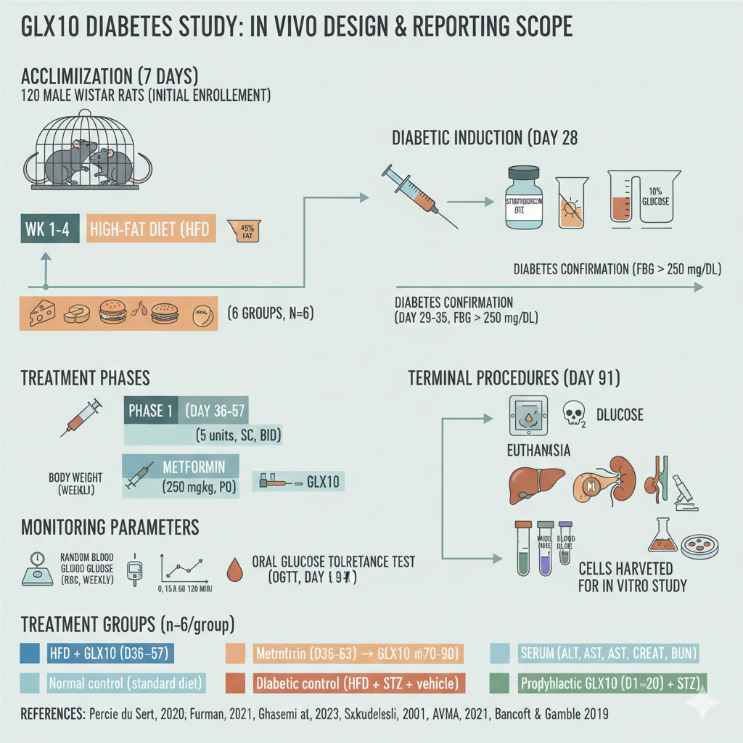

Supplementary S1: Experimental Design and Timeline of the In Vivo Study Assessing GLX10 in an HFD-STZ Rat Model of Type 2 Diabetes. The schematic outlines the transition from a broad 120-animal screening phase to a focused 6-group reporting phase (n=36). Following a 7-day acclimatization period, metabolic dysfunction was induced via a 4-week high-fat diet (HFD) followed by a low-dose streptozotocin (STZ) injection (25 mg/kg). The timeline details three distinct intervention strategies: prophylactic GLX10 (pre-STZ), acute GLX10 treatment (Phase I), and a sequential Metformin-to-GLX10 transition (Phase II). Longitudinal monitoring included weekly body weight and random blood glucose (RBG), culminating in an Oral Glucose Tolerance Test (OGTT) and terminal analysis of systemic (serum/CBC) and organ-specific (liver, pancreas, kidney) parameters.

Supplementary Figure S2 Body Weight Trends

Body Weight (Mean) over Days

|  | G1 HFD + GLX10 D36-57 | G2 Post oral + metf D36-63 + D70-D90 GLX10 | G3 HFD + Metformin Control | G4 Normal Control | G5 Diabetic Control | G6 Prophylactic GLX10 D1-D20 +stz |
| --- | --- | --- | --- | --- | --- | --- |
| 0 | 156.167 | 157 | 153 | 156 | 159.167 | 160 |
| 7 | 174.833 | 182 | 160.833 | 174 | 181.5 | 174.833 |
| 14 | 215.333 | 217.667 | 177.333 | 190.333 | 213 | 190.333 |
| 21 | 215.167 | 223.5 | 200 | 198.333 | 220.667 | 200.833 |
| 28 | 213.5 | 207.167 | 196.667 | 196.333 | 228.333 | 186.833 |
| 35 | 190.667 | 192.5 | 174.833 | 201.5 | 243.167 | 170 |
| 42 | 187.5 | 177.333 | 162.833 | 237.667 | 230.167 | 165.667 |
| 49 | 169.333 | 170.667 | 167.333 | 248 | 223.167 | 151 |
| 56 | 175.5 | 171.667 | 170 | 264 | 221.5 | 149.667 |
| 63 | 162.167 | 175.167 | 150.667 | 294.667 | 217.5 | 148 |
| 70 | 125 | 144.833 | 134.833 | 305.167 | 200.333 | 138.833 |
| 77 | 141 | 149.333 | 149.167 | 294 | 195.667 | 139 |
| 84 | 125.333 | 134.167 | 133 | 299.167 | 170.667 | 134.833 |
| 91 | 130 | 130.167 | 134.167 | 303.667 | 172.667 | 134 |

Supplementary S2: Longitudinal body weight changes for the six reported in-vivo groups across study days.

Supplementary Figure S3 Cell Viability SRB

| 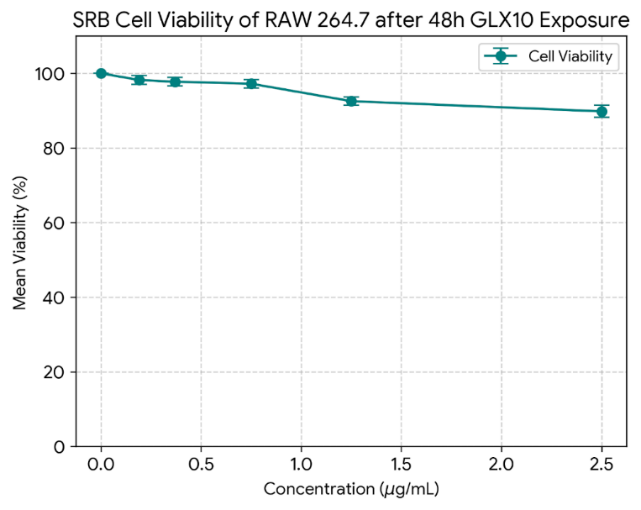 | RAW 264.7 macrophage viability remains relatively preserved across the tested range (0.19–2.5 µg/mL), supporting interpretation of NO inhibition under non-lethal exposure conditions. We can notice *High Viability:* At the highest concentration tested (2.50 µg/mL), the mean viability remained relatively high at approximately 89.8%, suggesting that the doses used in this range are well-tolerated by the RAW 264.7 macrophage cell line, *Minimal Cytotoxicity:* Between the control and 0.75 µg/mL, the viability drop was minimal (less than 3%), indicating negligible cytotoxicity at lower doses and the *Trend:* There is a clear downward trend in viability as the concentration increases from 0.75 µg/mL to 2.50 µg/mL. The above figures represent SRB-based viability dose–response curves of GLX10 in BNL (72 h), HepG2 (72 h), and RAW 264.7 (48 h). IC_50_ values were obtained from NAWAH curve fitting. GLX10 shows stronger cytotoxic potency against HepG2 carcinoma cells than BNL normal liver cells (IC_50_ HepG2 2.68 vs BNL 4.52 µg/mL), suggesting a cytotoxicity separation between malignant and normal hepatocyte models in this dataset. GLX10 also demonstrates a strong inhibitory profile against nitric oxide production in RAW264.7 (IC_50_ 0.98 µg/mL, see next figure). RAW 264.7 does not reach 50% loss of viability within the tested concentration range in the NAWAH RAW SRB dataset.” (Skehan et al., 1990). |
| --- | --- |

Supplementary S3: SRB viability in RAW 264.7

macrophages (48 h).

Supplementary Figure S4 Apoptosis AnnexinVPI.png

| 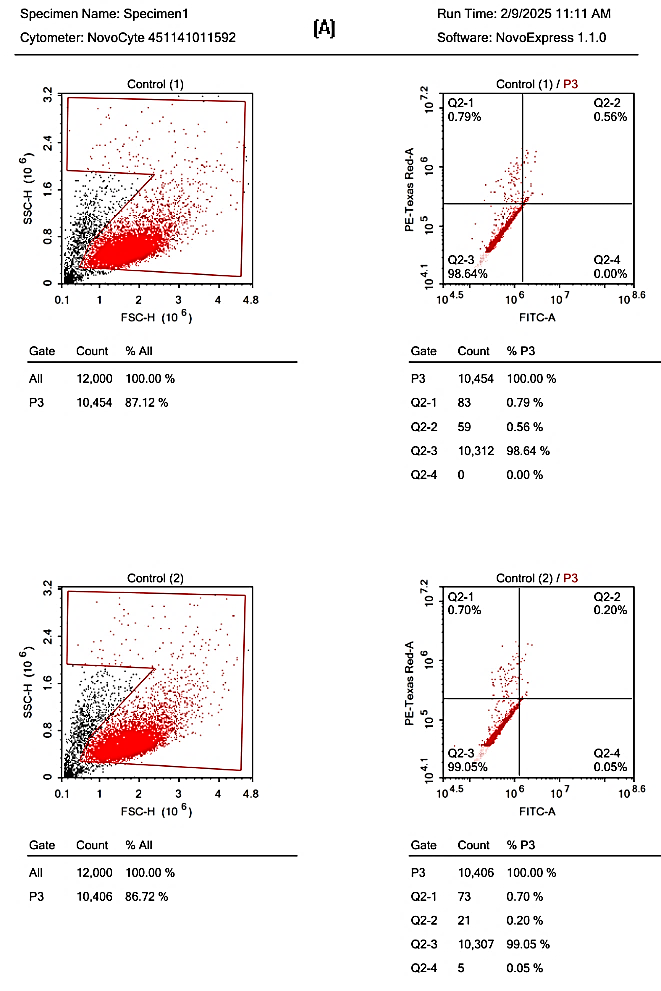 | 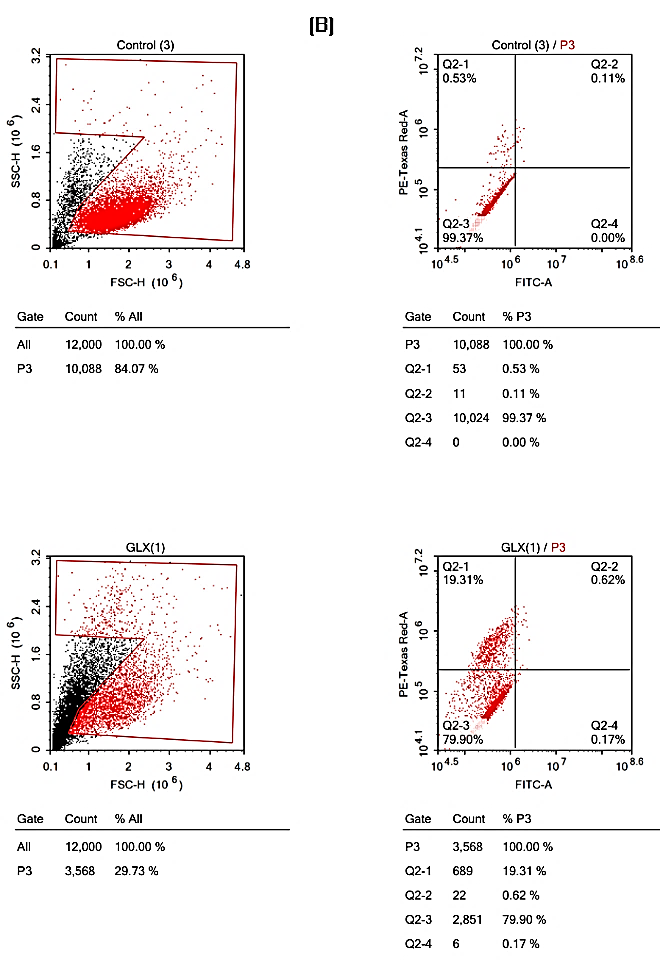 |
| --- | --- |
| 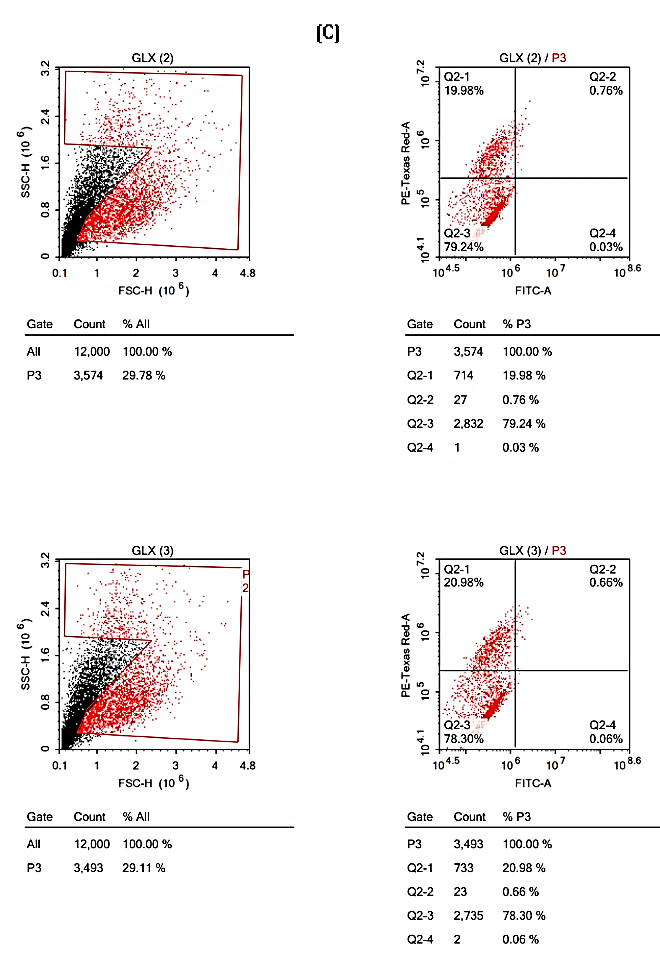 | Supplementary S4**: Annexin V/PI dot plots (BNLapoptosis)**   A. ***Control replicates:*** Representative Annexin V/PI dot plots for Control BNL cells (replicates), illustrating the dominant event distribution observed under control conditions. B. ***Control vs GLX10***: Representative dot plots comparing Control and GLX10-treated BNL cells, illustrating the redistribution of events across quadrants following GLX10 exposure. C. ***GLX10 replicates:*** Representative Annexin V/PI dot plots for GLX10-treated BNL cells (replicates), demonstrating consistency of the observed quadrant redistribution across replicates.   Autophagy-related readout (Acridine Orange staining, flow cytometry; BNL cells; 72 h) Autophagy-related AO flow readouts are summarized using the metrics reported by the NAWAH software output: Mean X (fluorescence intensity summary metric) and P1% (gated population percentage). These are reported exactly as provided in the annex output. |

Supplementary Figure S5 Autophagy Acridine Orange.png

| 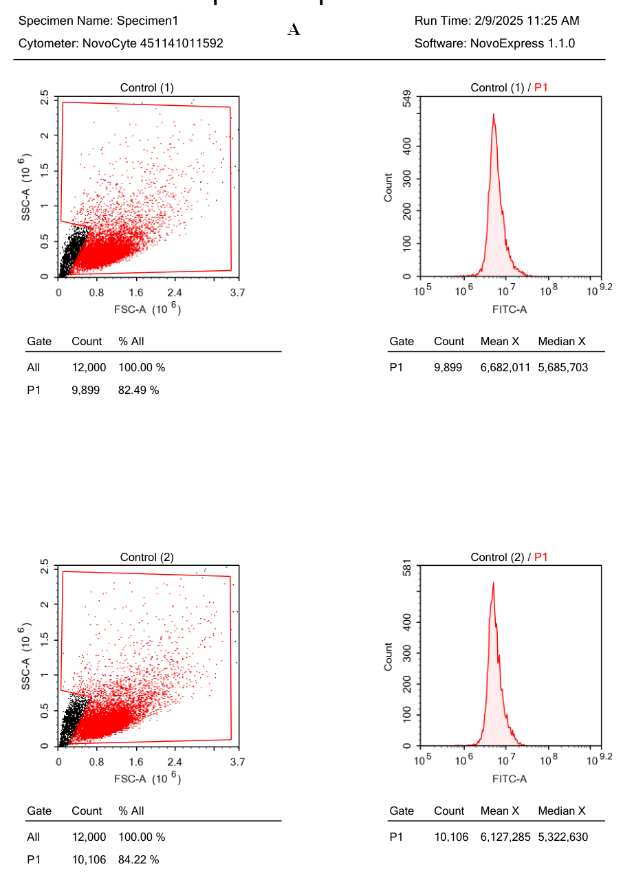 | 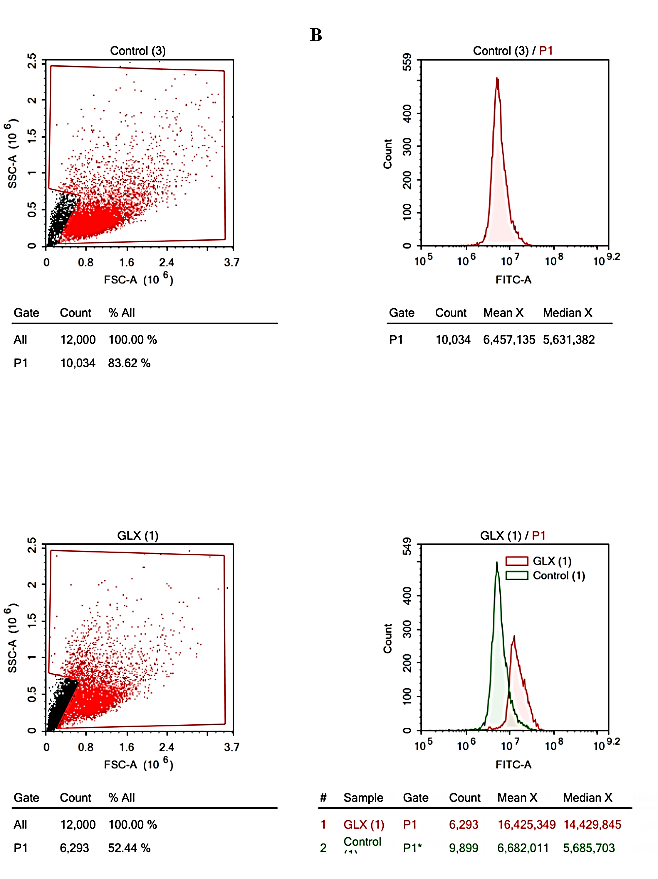 |
| --- | --- |
| 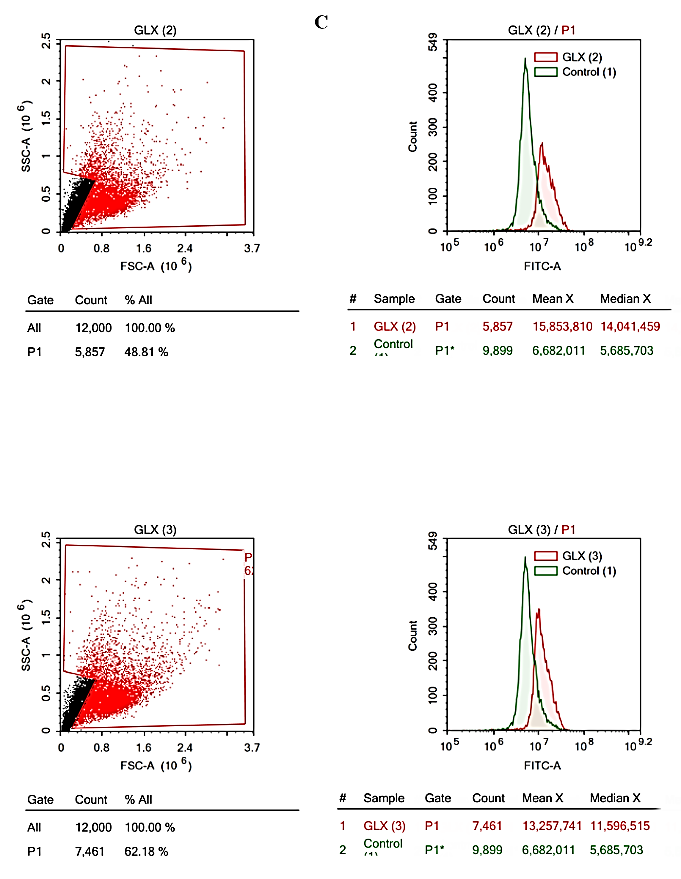 | **Supplementary S5 Acridine Orange flow-cytometry plots for Autophagy-related readout BNL cells (72 h).**  *The figures show the distribution shifts and fluorescence readouts reported for Control and GLX10 conditions (Thomé et al., 2016):*  *A. Control x Control.*  *B. Control x GLX10.*  *C. GLX10 x GLX10.*  **C**ell cycle profiling (DNA content flow cytometry; BNL cells; 72 h) Cell cycle phase fractions (G1, S, G2) and Sub-G1 fraction were extracted directly from the NAWAH annex outputs for three replicates per group (n = 3). Sub-G1 is reported as labeled by the software output. |

Supplementary Figure S6 Cell Cycle Analysis.png

| 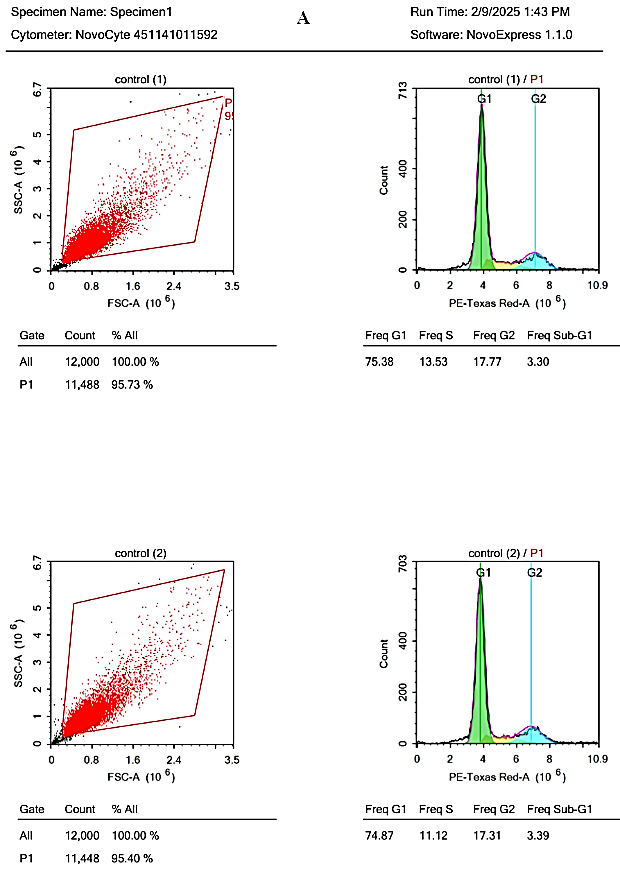 | 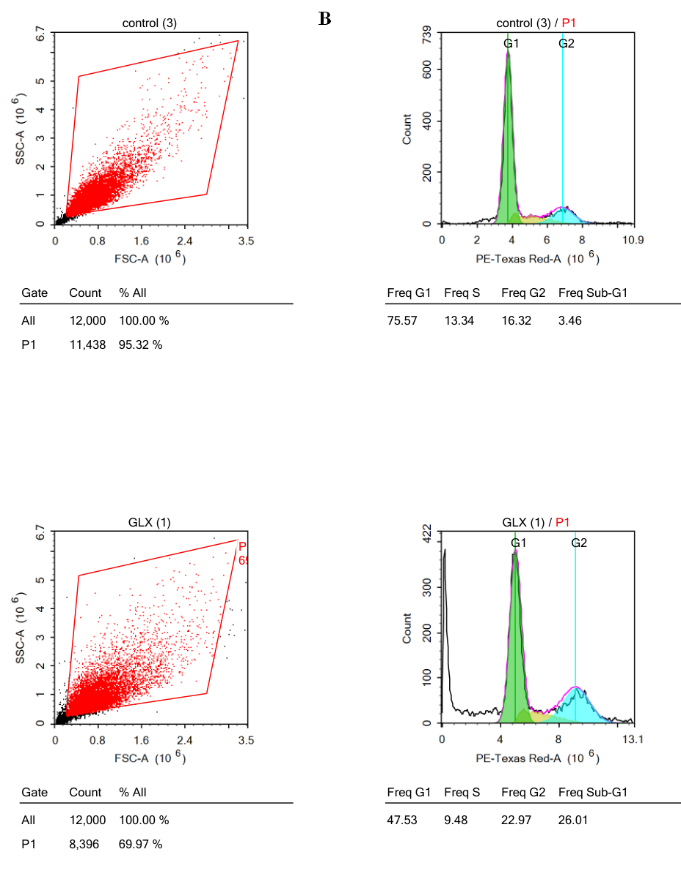 |
| --- | --- |
| 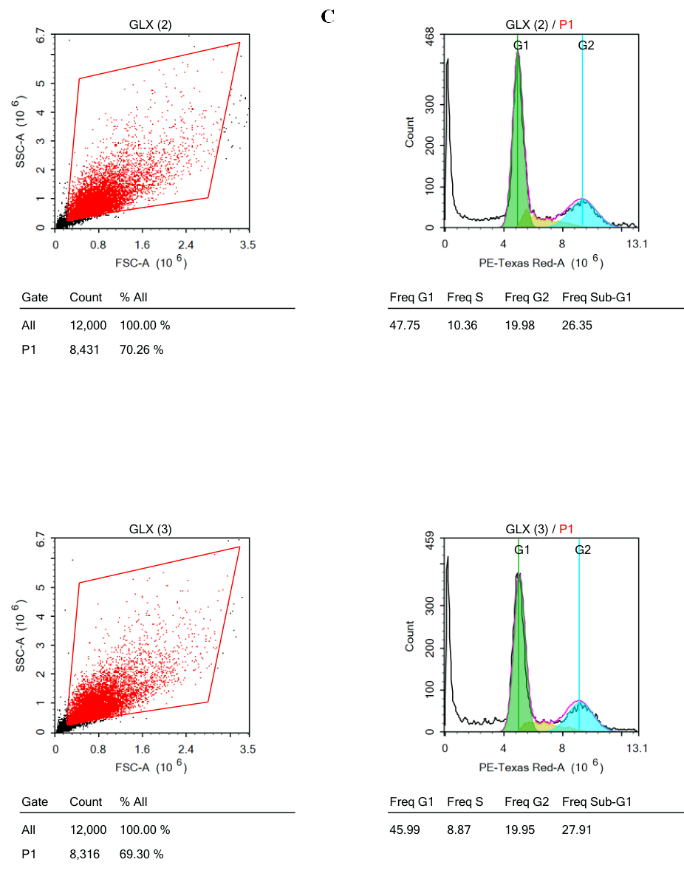 | **Supplementary S6: Cell-cycle flow-cytometry plots for BNL cells (72 h)**  *The figures show the flow-cytometry readouts under Control and GLX10 conditions, illustrating the phase distribution outputs (including Sub-G1):*  *A. Control x Control.*  *B. Control x GLX10.*  *C. GLX10 x GLX10.* |

Supplementary Figure S7 Western Blot Signaling.png

| 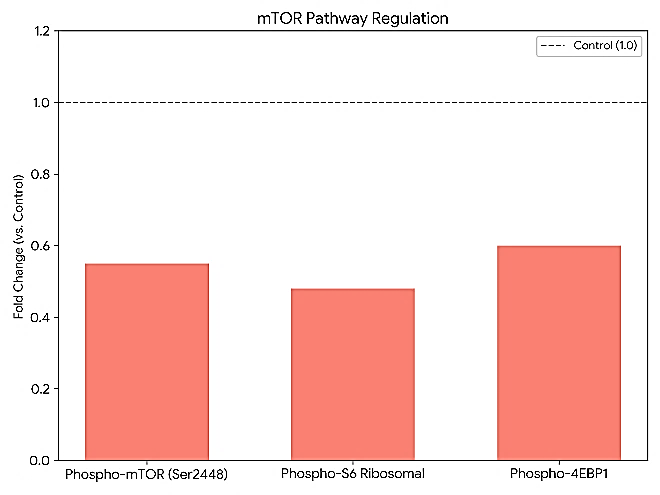 | 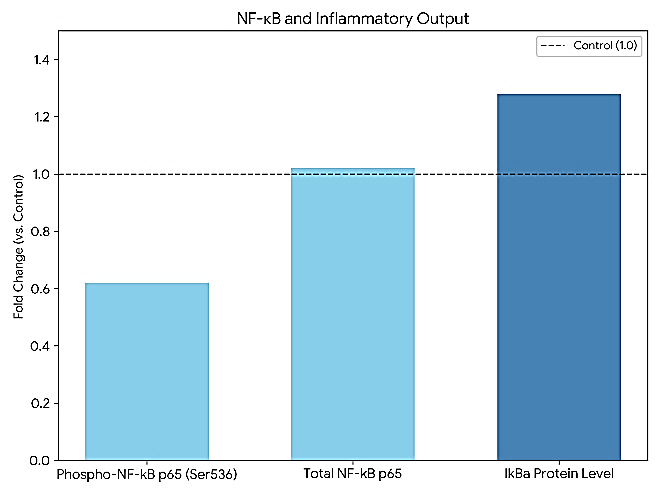 |
| --- | --- |
| 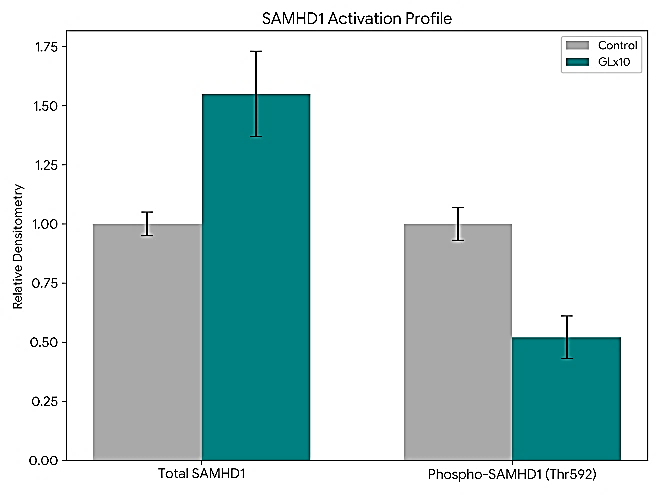 | Supplementary S7: Representative Western blot bands and densitometry-reported fold-change values relative to Control for SAMHD1, NF-κB pathway markers, and mTOR-axis phosphorylation markers in RAW 264.7 cells under GLX10 exposure (Liu et al., 2017; Wang & Proud, 2006).   - Upper Left: mTOR Pathway Inhibition: The markers for the mTOR pathway showed a consistent downregulation. Phospho-mTOR (Ser2448), Phospho-S6, and Phospho-4EBP1 were reduced to 0.55, 0.48, and 0.60-fold of control levels, respectively. This demonstrates that GLX10 effectively suppresses the nutrient-sensing mTOR signaling axis. - Upper Right: Anti-Inflammatory Response (NF-κB): GLX10 exerted a dual effect on the NF-κB pathway by reducing the activation of the pro-inflammatory subunit (Phospho-NF-κB p65 (Ser536) to 0.62-fold) and increasing the levels of the inhibitory protein IκBα to 1.28-fold. Total NF-κB p65 levels remained stable at 1.02-fold. - Lower Left: SAMHD1 Activation Profile: GLX10 treatment led to a significant increase in Total SAMHD1 protein levels (1.55-fold) while simultaneously reducing its Phosphorylation (Thr592) to 0.52-fold compared to the control. This combination indicates a strong activation of the SAMHD1 protein. |

Supplementary Figure S8 Mechanistic Model.png

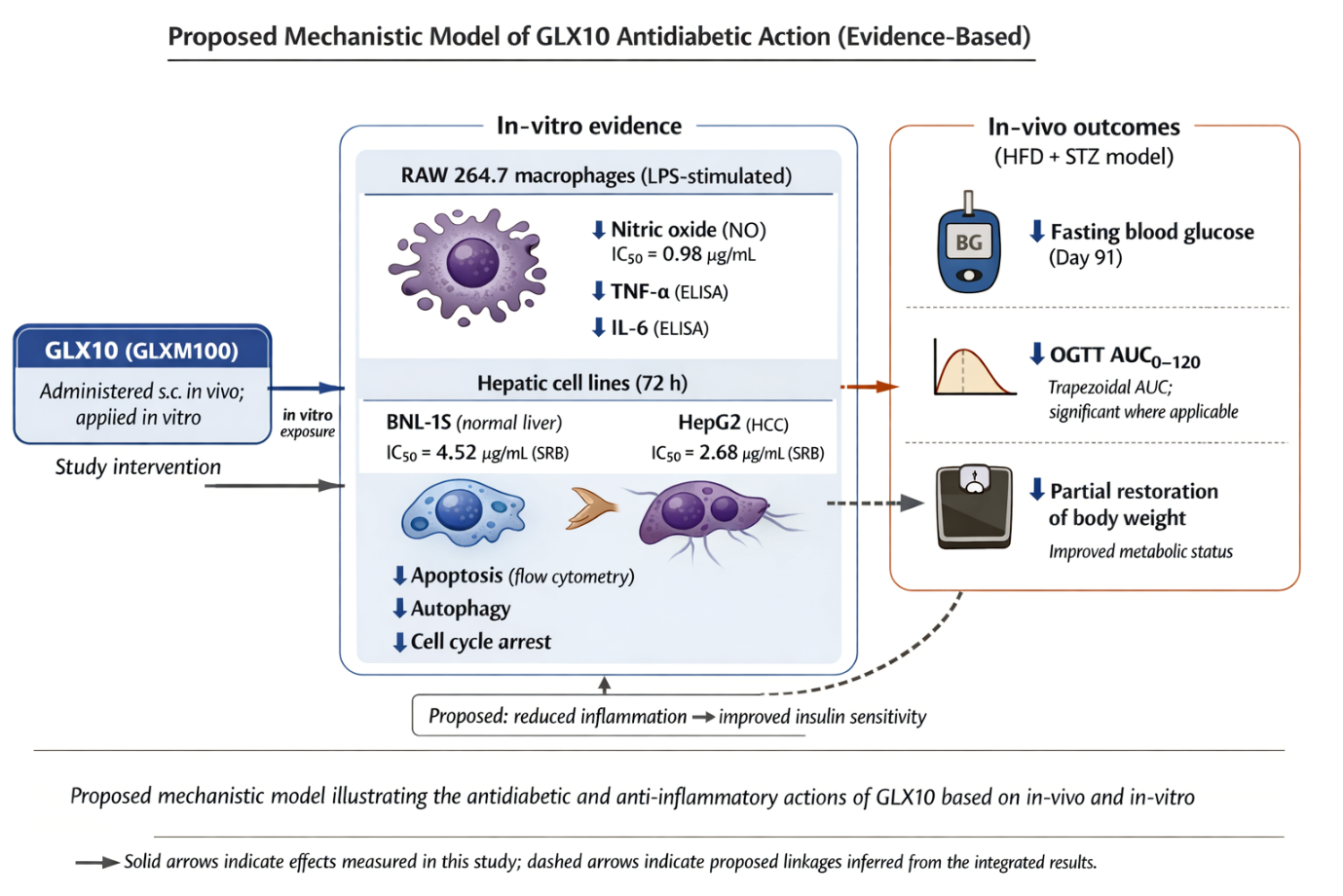

Supplementary S8: Proposed mechanistic model illustrating the antidiabetic and anti-inflammatory actions of GLX10 based on in-vivo and in-vitro findings.
